# D-Alanine esterification of teichoic acids contributes to *Lactobacillus plantarum* mediated intestinal peptidase expression and *Drosophila* growth promotion upon chronic undernutrition

**DOI:** 10.1101/098434

**Authors:** Renata C. Matos, Hugo Gervais, Pauline Joncour, Martin Schwarzer, Benjamin Gillet, Maria Elena Martino, Pascal Courtin, Sandrine Hughes, Marie-Pierre Chapot-Chartier, François Leulier

## Abstract

The microbial environment influence animal physiology. However, the underlying molecular mechanisms of such functional interactions are largely undefined. Previously, we showed that upon chronic undernutrition, strains of *Lactobacillus plantarum*, a dominant commensal partner of *Drosophila*, promote host juvenile growth and maturation partly via enhanced expression of intestinal peptidases. By screening a transposon insertion library of *Lactobacillus plantarum* in gnotobiotic *Drosophila* larvae, we identify a bacterial cell wall modifying machinery encoded by the *pbpX2-dltXABCD* operon that is critical to enhance host digestive capabilities and promote growth and maturation. Deletion of this operon leads to bacterial cell wall alteration with a complete loss of teichoic acids D-alanylation. We thus conclude that teichoic acids modifications participate in commensal-host interactions and specifically, D-alanine esterification of teichoic acids contributes to optimal *L. plantarum* mediated intestinal peptidase expression and *Drosophila* juvenile growth upon chronic undernutrition.

**Highlights:**

- *Lp^NC8^* mutant library screening identifies genes affecting *Drosophila* growth promotion.
- *pbpX2-dlt* operon is required for D-alanylation of teichoic acids and *Drosophila* growth.
- Deleting the *pbpX2-dlt* operon alters host intestinal peptidase expression.
- Peptidoglycan and *pbpX2-dlt* dependent signals are required for *Lp^NC8^* mediated growth promotion.

**eTOC blurb:** Animals establish interactions with their microbial communities that shape many aspects of their physiology including juvenile growth. However, the underlying molecular mechanisms are largely undefined. Matos *et al.* reveal that bacterial teichoic acids modifications contribute to host juvenile growth promotion.

## INTRODUCTION

Metazoans establish complex interactions with their resident microorganisms for mutual benefits (Hooper and Gordon, 2001). When in homeostasis, these interactions contribute to different aspects of host physiology (McFall-Ngai et al., 2013). In the gut, microbial communities enhance digestive efficiency by providing enzymatic functions that help their hosts optimize extraction of dietary energy and nutrients. In addition, the gut microbiota promotes proper immune system development, local immune homeostasis and limits pathogen colonization (Fraune and Bosch, 2010). Despite the renewed interest in understanding the functional impact of gut microbiota on host physiology, a clear view of the molecular dialogue engaged upon host/microbiota interaction remains elusive (Schroeder and Bäckhed, 2016). Therefore the use of simple animal models, such as *Drosophila*, may help unravel the evolutionarily conserved mechanisms underlying the impact of intestinal bacteria in their host physiology, since it combines genetic and experimental tractability with a cultivable microbiota and the ease to generate germ-free animals (Ma et al., 2015).

*Drosophila* gut microbiota is composed of simple and aerotolerant bacterial communities (mostly *Acetobacteraceae* and *Lactobacillaceae* families) with five prevalent species: *Acetobacter pomorum, A. tropicalis, Lactobacillus brevis, L. plantarum* and *L. fructivorans* (Wong et al., 2011). The genus *Lactobacillus* gathers bacteria with high phylogenetic and functional diversity (Goh and Klaenhammer, 2009). They have been largely used as model lactic acid bacteria and are recognized as potential health beneficial microorganisms in the human gastrointestinal tract (Kleerebezem et al., 2010). As a prevalent member of *Drosophila* microbiota, *L. plantarum* is involved in several aspects of host physiology such as social behaviour (Sharon et al., 2010), protection against infection (Blum et al., 2013), gut epithelial homeostasis (Jones et al., 2013), nutrition (Newell and Douglas, 2013; Wong et al., 2014) and post-embryonic development (Erkosar et al., 2015; Storelli et al., 2011). We previously reported that, upon chronic undernutrition, certain strains of *L. plantarum* (such as *L. plantarum^WJL^)* fully recapitulate the beneficial effect of a more complex microbiota by promoting *Drosophila* juvenile growth and maturation rate (Storelli et al., 2011). *L. plantarum^WJL^* exerts its beneficial effect on larval growth through the host nutrient sensing system that relies on the tissue specific activity of the TOR kinase, which subsequently modulates hormonal signals controlling growth and maturation (Storelli et al., 2011). Importantly, using conventional and gnotobiotic mice we recently demonstrated that the intestinal microbiota and some strains of *L. plantarum* also influence linear growth in mammals (Schwarzer et al., 2016). These results suggest that the still unknown molecular mechanisms underlying microbiota-mediated juvenile growth promotion are likely conserved during evolution. Recently, we showed that *L. plantarum* influences juvenile growth at least partly through the increased expression of a set of specific host digestive enzymes in the intestine (Erkosar et al., 2014; Erkosar et al., 2015). We have shown that *L. plantarum^WJL^* promotes the expression of peptidases, such as Jon66Ci and Jon66Cii, in the enterocytes in both PGRP-LE/Imd/Relish-dependent and independent manner. The resulting enhanced peptidase activity in the midgut increases the digestion of dietary proteins into dipeptides and amino acids as well as their uptake. Circulating dipeptides and amino acids are sensed in endocrine tissues by the TOR kinase pathway, which promotes the production of *Drosophila* growth factors, the insulin-like peptides (dILPs) and a precocious pic of production of the molting steroid hormone Ecdysone (Erkosar et al., 2015; Storelli et al., 2011).

Here, we aim to identify the bacterial genetic determinants involved in *L. plantarum-mediated* juvenile growth promotion and enhanced host digestive capabilities. Through the generation of a random transposon-mediated insertion library in the growth promoting strain *L. plantarum^NC8^*, we identified a set of bacterial genes whose function is critical to promote host growth. Among these genes, we further characterized the insertion in the *pbpX2-dltXABCD* operon with predicted functions in cell wall biogenesis and remodelling. Deletion of this operon alters the bacterial cell wall due to complete loss of teichoic acids D-alanylation, and significantly impairs *Drosophila* larval growth, maturation and intestinal peptidases expression. Our analysis points to the existence of additional host signalling pathway(s) besides the classic PGRP-LE/Imd/Relish involved in the sensing of the cell wall features defined by the gene products of *pbpX2-dltXABCD* operon. Taken together our results demonstrate that D-alanine esterification of teichoic acids contributes to *L.plantarum* mediated intestinal peptidase expression and *Drosophila* growth promotion upon chronic undernutrition.

## RESULTS

### Generation of a *L. plantarum^NC8^* random-mutagenesis library

To identify *L. plantarum (Lp)* genes required to sustain *Drosophila* juvenile growth and maturation, we adopted a classical unbiased forward genetic approach via transposon-mediated mutagenesis of the bacterial chromosome coupled to a phenotypical screening in mono-colonized animals. The previously characterized strain *Lp^WJL^* has low transformation efficiency (Matos R., unpublished results) and carries plasmids (Kim et al., 2013; Martino et al., 2015) that preclude random transposition in the bacterial chromosome. We therefore chose a *Lp* strain with high transformation efficiency, no native plasmids and capable to promote host growth in mono-association experiments to the same extent as *Lp^WJL^.* This strain, designated as *Lp^NC8^*, is suitable for transposon mutagenesis library construction (Gury et al., 2004). Upon mono-colonization both strains strongly support *Drosophila’s* larval linear growth (Fig. 1A) and maturation (i.e entry to metamorphosis; Fig. 1B) under chronic undernutrition when compared to the mildly growth promoting strain *Lp^NIZO2877^* (Schwarzer et al., 2016) or the germ-free (GF) condition (Fig. 1A-B). Thus, we mutagenized the *Lp^NC8^* chromosome using the P_junc_-TpaseIS_*1223*_ transposon mutagenesis system recently described by Licandro-Seraut et al. (2012). This system was developed for lactic acid bacteria and has been successfully applied to *Lactobacillus casei* (Licandro-Seraut et al., 2014) and *Lactobacillus pentosus* (Perpetuini et al., 2013). It couples a thermo-sensitive plasmid (pVI129) expressing transiently the IS_*1223*_ transposase and a suicide plasmid (pVI110) encoding the IS_*1223*_ transposon, which together lead to random insertion of IS_*1223*_ sequences into the bacterial chromosome. Strain NC8pVI129 (Table S2) was transformed with pVI110 and 2091 colonies were randomly selected and stocked as individual clones at −80°C. To evaluate the randomness of the transposon insertions in our library, we tracked transposon insertion sites by deep sequencing of flanking genomic sequences (see Methods). Sequencing reads were mapped to the *Lp^NC8^* genome, which revealed that transposon insertions are evenly distributed over *Lp^NC8^* genome with an average insertion site every 2kb (Fig. 1C). By analysing sequencing reads and insertions sites, we found that among the 2091 insertions, 1574 insertions disrupted 1218 different ORFs (42% of *Lp^NC8^* ORFs currently annotated; Table S1) and 517 landed in intergenic regions. Insertions were detected in ORFs belonging to different clusters of orthologous groups (COGs) (Table S1; Fig. S1) and globally, only minimal differences in the relative proportion of functional groups targeted in our library were observed as compared to the repartition of COGs in the *Lp^NC8^* genome (Fig. S1). These results demonstrate the insertion library sufficiently covers the genome in random manner, thus making it suitable for further phenotypic screen.

**Figure 1.**
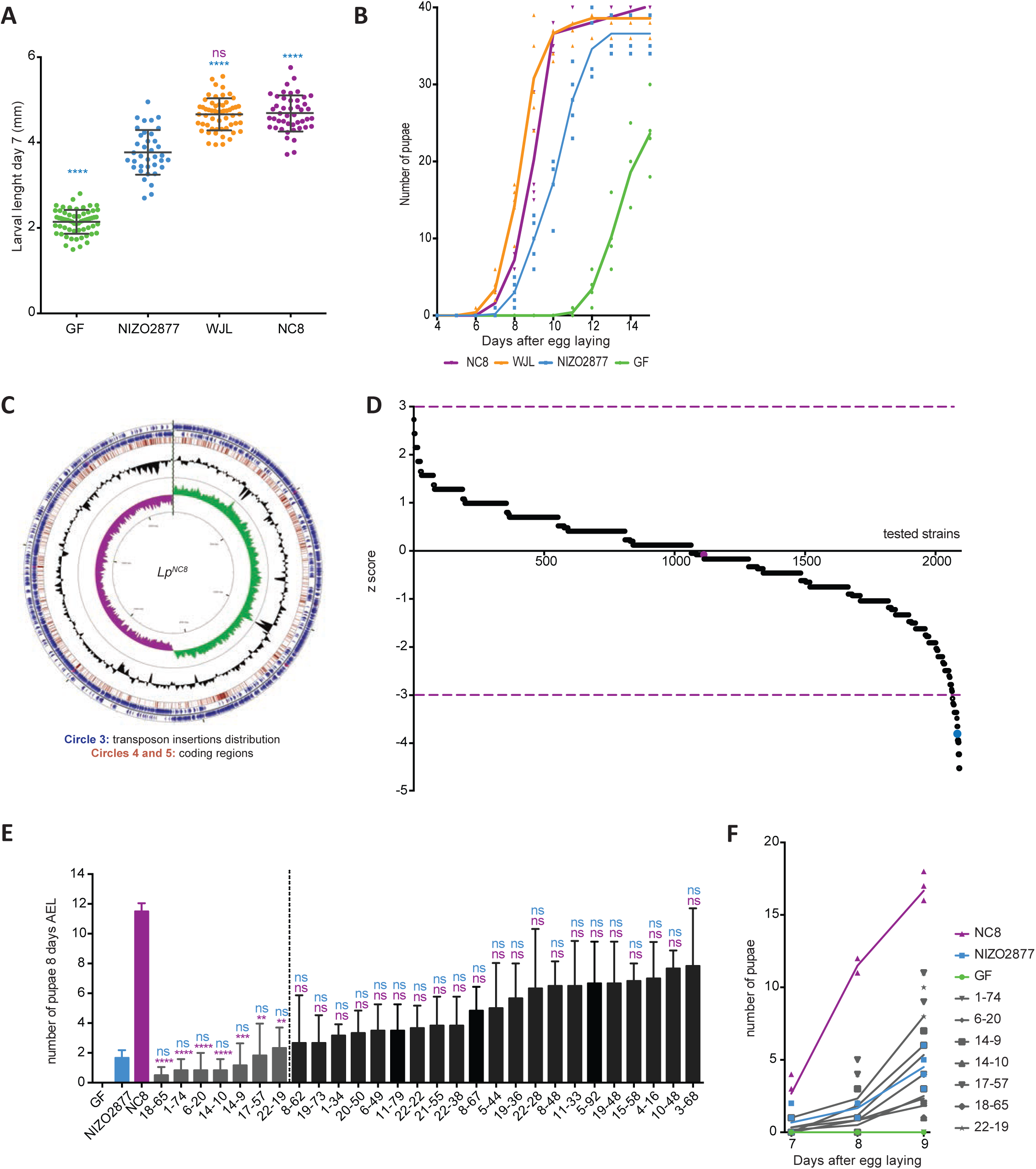
Identification of *Lp* genes involved in *Drosophila* growth promotion. (A) Larval longitudinal length after inoculation with *Lp^WJL^*, *Lp^NC8^* and *Lp^NIZO2877^* strains. Larvae were collected 7 days after association and measured as described in the Methods section. Blue asterisks illustrate statistically significant difference with larval size of *Lp^NIZO2877^; ns* represent absence of statistically significant difference between *Lp^WJL^* and *Lp^NC8^* larval sizes. ****: p<0,0001. (B) Number of emerged pupae scored over time from eggs associated with the strains *Lp^WJL^, Lp^NC8^, Lp^NIZO2877^* or PBS (for the GF condition). Forty GF eggs were associated independently with 10^8^ CFUs of each one of the strains in 5 replicates. The number of pupae was scored every 24 h. (C) Genome atlas of transposon insertions mapped to *Lp^NC8^* genome. Visualization of the transposon insertions mapped onto the *L. plantarum^NC8^* genome. The innermost rings represent the GC skew (Circle 1) in purple/green and GC content (Circle 2) in black. Circle 3 shows the distribution of the genomic regions disrupted by each transposon insertion (red bars). Circle 4 and 5 show *Lp^NC8^* coding regions (blue arrows), tRNAs (red arrows) and rRNAs (purple arrows) on the negative and positive strand respectively. (D) Screen of the random transposon insertion library for mutants with altered growth promotion phenotype under chronic under-nutrition. The number of pupae scored at day 8 after association with the 2091 insertional mutants were converted in z-scores. Purple lines indicate the cut off of z=+/−3. Control strains are represented by colored dots: *Lp^NC8^* in purple and *Lp^NIZO2877^* in blue. (E) Number of emerged pupae 8 days after association of GF eggs with the 28 loss-of-function candidates selected form the primary screen after setting the z score at −3. Purple asterisks illustrate statistically significant difference with *Lp^NC8^* number of pupae; *ns* represent absence of statistically significant difference with *Lp^NC8^* (purple) and *Lp^NIZO2877^* (blue). ****: p<0,0001; ***: 0,0001<p<0,001; **: 0,001<p<0,01. The candidate strains on the left of the dotted line were retained for further analyses. (F) Number of emerged pupae at days 7, 8 and 9 after association of 20 GF eggs with 10^8^ CFUs of the 7 loss-of-function candidates selected from the secondary screen.

### Screening of the *Lp^NC8^* mutant library identifies mutants affecting *Lp*-mediated larval growth promotion

We screened the insertions library with the aim to identify *Lp^NC8^* mutants, which upon mono-colonization have an altered capacity to sustain growth and maturation of chronically undernourished *Drosophila.* Prior to the library screening, we experimentally determined that in these nutritional conditions, the most robust timing to visually discriminate (by counting the number of pupae emerging from food) a strong growth promoting strain *(Lp^NC8^)*, a moderate growth promoting strain (*Lp^NIZO2877^*; Schwarzer et al., 2016)) and the GF condition is day 8 after GF eggs inoculation with bacterial isolates (Fig. 1B, S2). We searched for *Lp^NC8-pVI110^* insertion mutants with a growth promotion capacity weaker or similar to *Lp^NIZO2877^* (Fig. 1B, S2). The screen was conducted as follow: 20 GF eggs were deposited in tubes containing low-yeast diet which were inoculated independently with each one of the 2091 *Lp^NC8-pVI110^* insertion strains, while *Lp^NIZO2877^*, *Lp^NC8^* and GF served as controls. After 8 days of development, the number of emerging pupae was scored in each tube and normalized into z scores (i.e the score for each strain reflecting the number of standard deviations from the mean number of pupae scored at day 8 from all tested strains; mean=15,18 pupae, SD=3,53; Fig. 1D). The control strains yielded z scores of −0,05 for *Lp^NC8^* and −3,73 for *Lp^NIZO2877^*. This screen revealed insertions leading to either gain or loss of function phenotypes as compared to the reference strain *Lp^NC8^* with z scores ranging from +2,73 down to - 4,52 (Fig. 1D). In order to select robust loss-of-function candidates similar or stronger than the *Lp^NIZO2877^* strain, we set a selection cut-off for z scores below −3 and identified 28 transposon insertion mutants, which were retained for a secondary screening procedure. These candidates were re-tested in 3 independent experiments as during the primary screen but this time the number of emerging pupae was recorded at days 7, 8 and 9. From the secondary screen, we confirmed that inoculations with 7 insertion mutants robustly delayed pupariation time with statistical significance when compared to the wild-type (WT) *Lp^NC8^* strain and were statistically undistinguishable from the *Lp^NIZO2877^* strain (Fig. 1E-F). To exclude the possibility that the loss of the growth promoting phenotype of these 7 mutants was a consequence of their inability to persist in the low yeast fly food, we assessed bacterial loads in the food matrix at days 3, 5, 7 and 10 after initial inoculation of 10^8^ CFUs.mL^−1^ for each strain (Fig. S3). All the mutant strains behave as the *Lp^NC8^* strain at each time point.

Having characterized the impact of the 7 insertion mutants on *Drosophila* growth promotion, we identified the transposon insertion sites in *Lp^NC8^* genome. Of the 7 insertions, 6 were inside ORFs and 1 in an intergenic region between the ORFs encoding the transporter *secE* and the transcription regulator *nusG* (Table 1 and Fig. S4). The insertions in ORFs hit the *mleP1* gene encoding a malate transport protein (referred to as 1-74), the *pbpX2* gene encoding a putative serine-type D-D-carboxypeptidase (referred to as 6-20), the *pts28ABC* gene encoding a PTS system component (referred to as 14-10), the *lp_2466* gene encoding a prophage terminase large subunit (referred to as 17-57), the *lp_1944* gene encoding an ABC transporter (referred to as 18-65) and the *dnaK* gene encoding a chaperone protein (referred to as 22-19) (Table 1). Previously, we reported that peptidoglycan recognition by PGRP-LE contributes to *Lp^WJL^*-mediated intestinal peptidase enhanced expression during juvenile growth promotion (Erkosar et al., 2015). Hence, we were intrigued by the 6-20 mutant, hitting *pbpX2*, with a predicted function in peptidoglycan biosynthesis/maturation (Neuhaus and Baddiley, 2003; Palumbo et al. 2006), we thus pursued the characterization of 6-20 mutant.

**Table 1.** Genes disrupted by the transposon in the loss-of-function candidates.

| Mutant number | Locus of pVI110 disruption | Annotation |
| --- | --- | --- |
| 1-74 | *lp_0594* | Malate transport protein (*mlep1*) |
| 6-20 | *lp_2021* | Serine-type D-ala-D-ala carboxypeptidase (*pbpX2*) |
| 14-9 | *IG lp0616-lp0617* | Intergenic region between *secE* and *nusG* |
| 14-10 | *lp_3240* | PTS system, beta-glucosides-specific EIIABC component (*pts28ABC*) |
| 17-57 | *lp_2466* | Prophage P2b protein 15, terminase large subunit |
| 18-65 | *lp_1944* | Multidrug ABC transporter |
| 22-19 | *lp_2027* | Chaperone, heat shock protein DnaK |

### Deletion of *pbpX2-dltXABCD* operon affects *Lp^NC8^* mediated larval growth promotion

To confirm the loss-of-function phenotype observed with the transposon insertion mutant 6-20, we generated a deletion mutant of the *pbpX2* gene in the *Lp^NC8^* strain by homology-based recombination (Δ*pbpX2;* Fig. 2A). We tested the ability of the newly constructed Δ*pbpX2* strain to promote *Drosophila’s* growth by determining larval size seven days post-inoculation (Fig. 2B). We also assessed the developmental timing (Fig. 2C) of GF eggs monoassociated with either *ΔpbpX2* or the *Lp^NC8^* (WT) strain. Larvae associated with *ΔpbpX2* strain are significantly smaller than those associated with *Lp^NC8^* and their larval development is significantly delayed (Fig. 2B-D). These results confirm the importance of *pbpX2* gene to *Lp^NC8^* growth promoting effect. However the effect observed in Δ*pbpX2* strain is not as pronounced as the one observed with the 6-20 insertion mutant (Fig. 2B-D). The *pbpX2* gene is located in the *dlt* locus and is the first ORF of an operon that encodes 5 additional genes located in 3’ position to *pbpX2*, the *dlt* genes (*dltXABCD*; Fig. 2A; Palumbo et al., 2006). Given that *pbpX2* is co-transcribed with *dltXABCD* genes (Palumbo et al., 2006), we wondered if the 6-20 insertion in the *Lp^NC8^* strain would generate a polar effect and disrupt the entire operon and not only the *pbpX2* gene. To test our hypothesis, we engineered a strain deleted for the entire *pbpX2-dltXABCD* operon *(Δdlt_op_)* by homology-based recombination (Fig. 2A). We tested the *Δdlt_op_* mutant in larval size and developmental timing assays and observed that *Δdlt_op_* now recapitulated the 6-20 mutant phenotype and showed an increased loss of function phenotype as compared to *ΔpbpX2* (Fig. 2B-D). Thus, we conclude that the 6-20 insertion in the *Lp^NC8^* strain disrupts the entire *pbpX2-dltXABCD* operon (and not just *pbpX2)*, which encodes important functionalities in *Lp^NC8^* to promote *Drosophila’s* growth.

**Figure 2.**
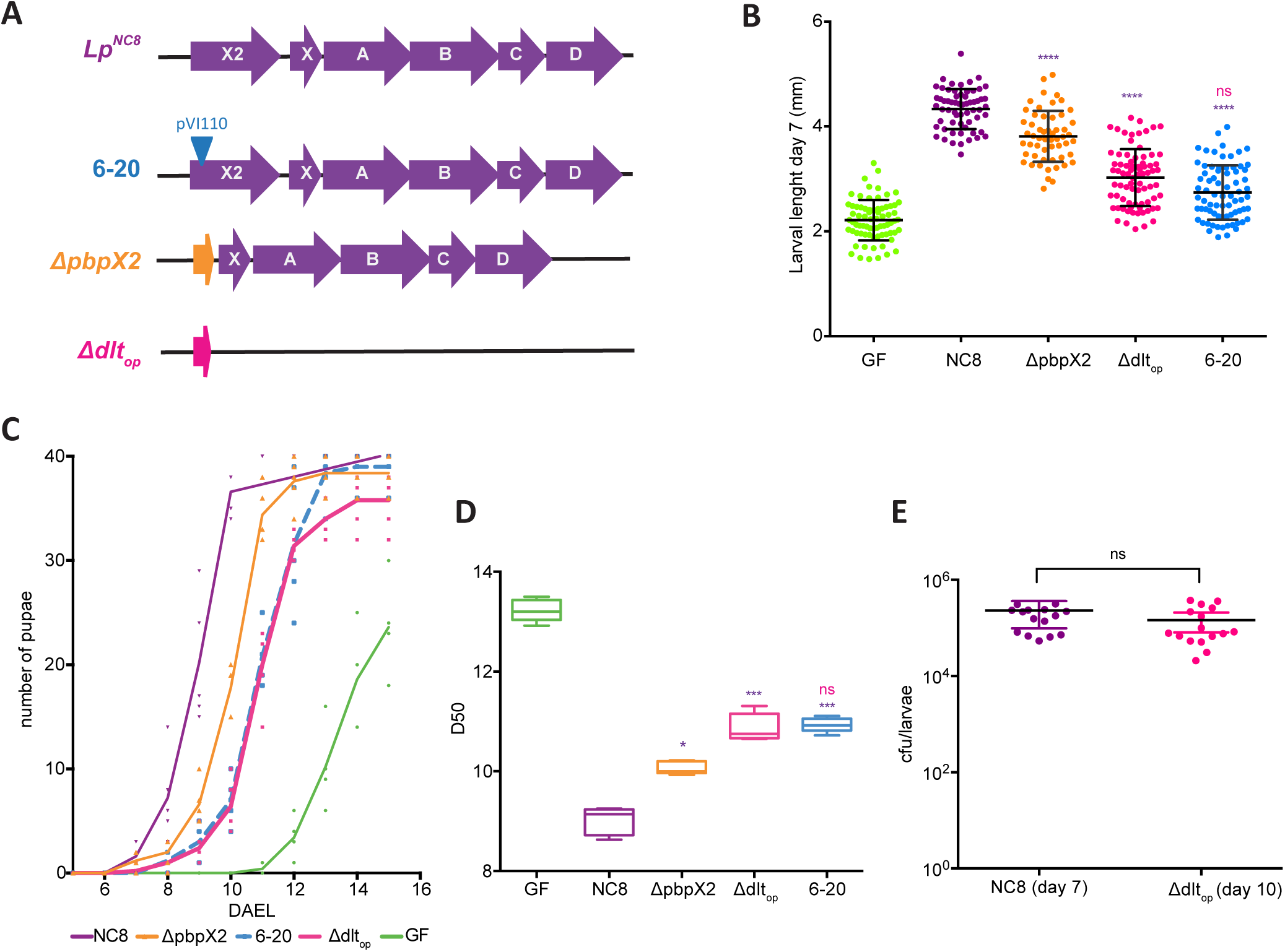
The *pbpX2-dltXABCD* operon impacts *Drosophila’s* growth. (A) *pbpX2-dlt* operon genomic organization in the *Lp^NC8^* strain: *pbpX2/dltX/dltA/dltB/dltC/dltD.* 6-20, pVI110 insertion within *pbpX2* gene. *ΔpbpX2* mutant, in which *pbpX2* gene was deleted by homology-based recombination. *Δdlt_op_* mutant, with genes *pbpX2/dltX/dltA/dltB/dltC/dltD* deleted by homology-based recombination. (B) Larval longitudinal length after inoculation with strains *Lp^NC8^*, 6-20, *ΔpbpX2, Δdlt_op_* or PBS (for the GF condition). Larvae were collected 7 days after association and measured as described in the Methods section. Purple asterisks illustrate statistically significant difference with *Lp^NC8^* larval size; *ns* represent absence of statistically significant difference with *Δdlt_op_*. ****: p<0,0001. (C) Number of emerged pupae scored over time for eggs associated with strains Lp, 6-20, ΔpbpX2, Δdlt_op_ or PBS (for the GF condition). Forty germ-free eggs were associated independently with 10^8^ CFUs of each one of the strains in 5 replicates. The number of pupae was scored every 24h. (D) Day when fifty percent of pupae emerge during a developmental experiment (D50) for GF eggs associated with strains *Lp^NC8^*, 6-20, *ΔpbpX2, Δdlt_op_* or PBS (for the GF condition). Purple asterisks illustrate statistically significant difference with *Lp^NC8^* larval size; ns represent absence of statistically significant difference with *Δdlt_op_*. ***: 0,0001<p<0,001; **: 0,001<p<0,01; *: p<0,05. (E) Bacterial load of size matched larvae associated with 10^8^ CFUs of *Lp^NC8^* (larvae collected 7 days after association) or *Δdlt_op_* (larvae collected 10 days after association). *ns* represents absence of statistically significant difference between the two conditions.

We next wondered if an altered colonization of the larvae by the mutant bacteria caused the *Δdlt_op_* loss of function phenotype. To test this hypothesis we quantified the load of *Lp^NC8^* and mutant bacteria associated to surface-sterilized and size-matched larvae. We did not detect a significant difference between the ability of *Lp^NC8^* and *Δdlt_op_* cells to associate with the larvae (Fig. 2E). This result indicates that the altered larval growth promotion of the *Δdlt_op_* mutant is not caused by a general disruption of the association between *Drosophila* larvae and its symbiont but probably by a rupture of the molecular dialogue engaged between those symbiotic partners.

### Deletion of *pbpX2-dltXABCD* genes in *Lp^NC8^* impacts cell morphology, D-alanylation of TA but not bacterial growth dynamics

Previously, inactivation of the operon encoding *dlt* genes in the strain *Lactobacillus plantarum^WCFS1^ (Lp^WCFS1^)* and the strain *Lactobacillus rhamnosus^GG^ (Lr^GG^)* had been associated with a major reduction of esterification of lipoteichoic acids (LTA) by D-alanines, reduced bacterial growth rate and increased cell lysis (Palumbo et al., 2006; Perea Vélez et al., 2007). Therefore we investigated *Lp^NC8^* and *Δdlt_op_* mutant growth dynamics by following optical density (at 600 nm) and CFU counts in liquid cultures (Fig. 3A). Similarly to the *dltB* mutant in the *Lp^WCSF1^* strain (Palumbo et al., 2006) *Δdlt_op_* mutant enters stationary phase with lower OD_600_ reading. However, while studying CFU counts, we did not detect any significant differences in bacterial cells doubling time (Fig. 3A; 98 minutes for *Lp^NC8^ vs* 94 minutes for *Δdlt_op_).* This is in line with our previous results posing that the 6-20 mutant bacteria persist in the low-yeast fly food similarly to wild-type *Lp^NC8^* cells (Fig. S3) and that *Δdlt_op_* mutant cells colonize larvae like *Lp^NC8^* cells (Fig. 2E). This striking difference between CFU counts and absorbance at 600nm for the two strains prompted us to study the cell morphology of *Lp^NC8^* and *Δdlt_op_* strains by phase contrast and fluorescence microscopy following membrane staining with FM4-64 (Fig. 3B). We detected a significant reduction of the width of *Δdlt_op_* mutant cells (Fig. 3B-C; *Lp^NC8^* mean width of 783 nm ±105; *Δdlt_op_* mean width of 658 nm ±104). These observations are reminiscent of the *dlt* genes loss of function phenotypes seen in *Lp^WCFS1^* and *Lr^GG^*, yet, in contrast to those strains, the change in cell shape of *Lp^NC8-Δdltop^* mutant is not associated with reduced cell viability.

**Figure 3.**
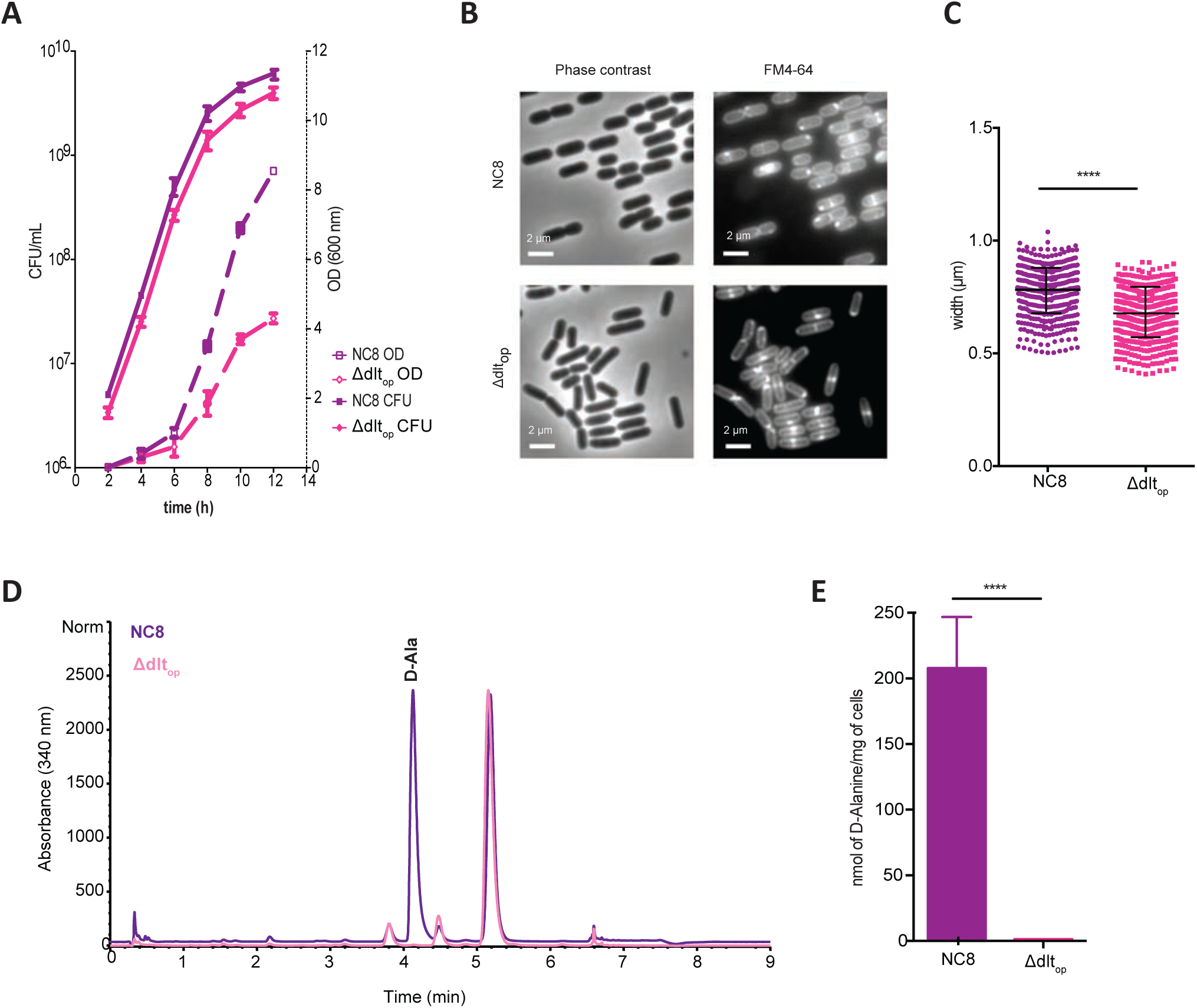
Cell envelope changes related to *pbpX2-dltXABCD* operon deletion. (A) Growth curves of *Lp^NC8^* and *Δdlt_op_* strains in MRS medium. Tubes containing sterile MRS medium were inoculated with 10^6^ CFUs of each strain and their growth was followed by optical density (OD) measurement at 600 nm and CFU counts by plating cultures dilutions in solid MRS. (B) Observation of *Lp^NC8^* and *Δdlt_op_* cells by phase contrast microscopy following membrane staining with FM4-64. Representative images from one out of three independent experiments. (C) *Lp^NC8^* and *Δdltop* cell width measurements from phase contrast microscopy observations. (D) HPLC detection of D-alanine released from whole cells. D-alanine released by alkaline hydrolysis from *Lp^NC8^* and *Δdlt_op_* whole cells were derivatized with Marfey’s reagent and separated by HPLC. D-alanine derivatives eluted at a retention time of 4.2 minutes. The eluted compounds were detected by UV absorbance at 340 nm. Quantification was achieved by comparison with D-ala standards in the range of 100 to 1500 pmol. (E) Amount of D-alanine released from whole cells of *Lp^NC8^* and *Δdlt_op_* by alkaline hydrolysis and quantified by HPLC. Mean values were obtained from five independent cultures with two injections for each.

Next, we determined the amount of D-alanine esterified to teichoic acids of *Lp^NC8^* and *Δdlt_op_* strains. D-alanine was released from dried bacteria by mild alkaline hydrolysis and was quantified by HPLC (Fig. 3D) as described previously (Kovács et al., 2006), allowing to estimate D-alanine esterified to both lipoteichoic acids (LTA) and wall teichoic acids (WTA). D-alanine was released in appreciable amounts from *Lp^NC8^* cells but was almost undetectable from the *Δdlt_op_* cells (Fig. 3E). We therefore conclude that the deletion of *pbpX2-dltXABCD* genes leads to a loss of D-alanine esterification on teichoic acids and size reduction in the mutant cells. Thus, these observations indicate that the enzymatic activity leading to D-alanine esterification of teichoic acids (TA) is required for *Lp^NC8^* mediated *Drosophila* growth promotion upon chronic undernutrition.

### Deletion of *pbpX2-dltXABCD* genes impacts intestinal peptidase expression

We previously reported that upon chronic undernutrition, *Lp* promotes *Drosophila* larval growth partly by increasing the expression of a set of intestinal peptidases (Erkosar et al., 2015). Elevated proteolytic activity in the intestine optimizes the digestion of dietary proteins into dipeptides and aminoacids (AAs), which, when taken up by cells, activate the TOR kinase pathway in the endocrine tissues to release dILPs and ecdysone, whose actions prompt systemic growth and maturation (Erkosar et al., 2015; Storelli et al., 2011). The activity of the PGRP-LE/Imd/Relish signalling cascade in the enterocytes regulates the expression of some intestinal peptidases in the presence of *Lp*, but loss of function mutations in the PGRP-LE/Imd/Relish cascade only partially dampen peptidase induction upon mono-association (Erkosar et al., 2015). We thus postulated that in addition to PG sensing and PGRP-LE/Imd/Relish signalling, additional compounds produced by the bacteria would impact intestinal peptidase expression and activity through yet unknown mechanisms. Given that *Δdlt_op_* strain is a poor *Drosophila* growth promoting strain with an altered cell wall composition, we hypothesize that the *pbpX2-dlt* dependent D-alanine esterification of TA might be the source of additional bacterial signals sensed by enterocytes to trigger intestinal peptidase expression.

To test this hypothesis, we investigated how the *Δdlt_op_* strain influences intestinal peptidases expression. To this end, we inoculated GF eggs with either sterile PBS, *Lp^NC8^* or *Δdlt_op_* strains, collected size matched larvae (Size 1 and 2, S1-S2) at an early and late larval stage (early L2 - S1 and mid-L3 - S2 respectively; Fig. 4A) and assayed intestinal peptidase expression. In control larvae (left part of each panel), the *Δdlt_op_* strain sustains *Jon66Ci* and *Jon66Cii* expression at lower levels than the *Lp^NC8^* strain - this tendency only becomes statistically significant for the *Jon66Cii* peptidase at mid-L3 stage (S2; Fig. 4B-C). This suggests that a host-sensing mechanism triggered by bacteria bearing D-alanylated TA in their cell wall indeed increases the expression of gut proteases.

**Figure 4.**
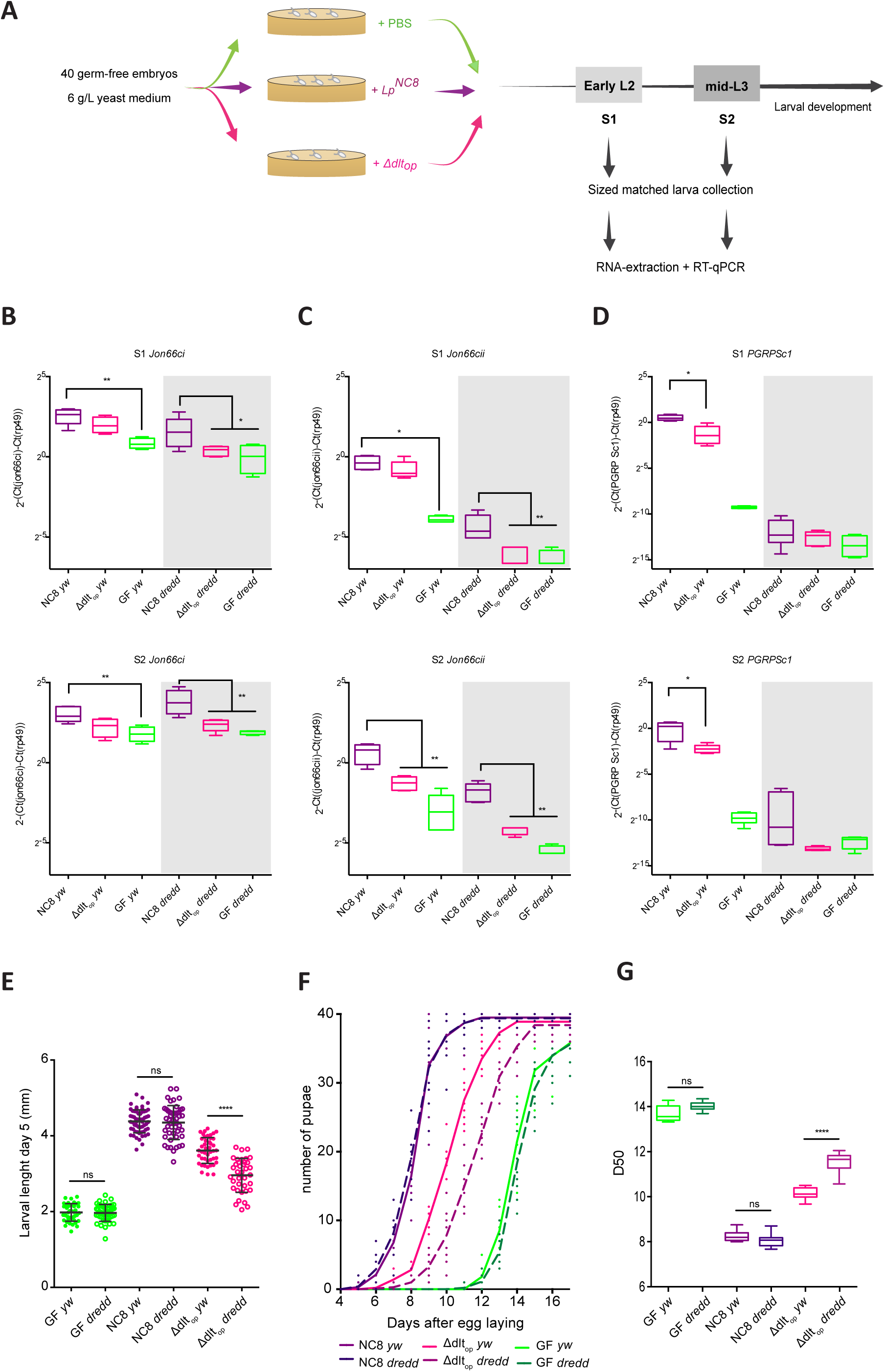
*Drosophila* reduced proteases expression in presence of *Δdlt_op_* strain is independent of IMD pathway. (A) Experimental set-up for the RT-qPCR analysis. Forty germ-free eggs of *y,w* and *y,w,Dredd* flies were inoculated with 10^8^ CFUs of *Lp^NC8^* or *Δdlt_op_* strains. For each condition, larvae were collected at early L2 and mid-L3 phases. Their RNA was extracted and RT-qPCR performed as described in the Methods section. (B) Mean ± SEM of ΔCt^gene^/ΔCt^rp49^ ratios for *Jon66Ci* detected in size matched larvae associated with *Lp^NC8^, Δdlt_op_* or the GF condition. **: 0,001<p<0,01; *: p<0,05. (C) Mean ± SEM of ΔCt^gene^/ΔCt^rp49^ ratios for *Jon66Cii* detected in size matched larvae associated with *Lp^NC8^, Δdlt_op_* or the GF condition. **: 0,001<p<0,01; *: p<0,05. (D) Mean ± SEM of ΔCt^gene^/ΔCt^rp49^ ratios for *PGRP-Sc1* detected in size matched larvae associated with *Lp^NC8^, Δdlt_op_* or the GF condition. *: p<0,05. (E) *y,w* and *y,w,Dredd* larval longitudinal length after inoculation with 10^9^ CFUs of *Lp^NC8^, Δdlt_op_* and PBS, for the germ-free condition. Larvae were collected 5 days after association and measured as described in the methods section. *ns* represents absence of statistically significant difference; ****: p<0,0001. (F) Number of *y,w* and *y,w,Dredd* emerged pupae scored over time for eggs associated with strains *Lp^NC8^, Δdlt_op_* and PBS for the germ-free condition. Forty germ-free eggs were associated independently with 10^9^ CFUs of each one of the strains in 5 replicates. The number of pupae was scored every 24h. (G) Day when fifty percent of pupae emerge during a developmental experiment (D50) for *y,w* and *y,w,Dredd* GF eggs associated with strains *Lp^NC8^, Δdlt_op_* and PBS (for the GF condition). *ns* represents absence of statistically significant difference; ****: p<0,0001.

### Sensing of both PG and *pbpX2-dltXABCD* dependent bacterial signals is required for *Lp^NC8^* mediated intestinal peptidase expression and larval growth promotion

Next, we tested if the impaired induction of gut peptidases by *Δdlt_op_* mutant is further altered in *Dredd* larvae. *Dredd* encodes an essential positive regulator of the PG-sensing Imd/Relish cascade (Leulier et al., 2003; 2000). In *Dredd* larvae, the PGRP-LE/Imd/Relish signalling cascade is completely inactivated in the enterocytes upon *Lp* association (Erkosar et al., 2015). We confirmed this observation by analysing the expression of *PGRP-SC1*, whose expression in the gut entirely relies on the PGRP-LE/Imd/Relish cascade upon *Lp* association (Erkosar et al., 2015). We found that while *PGRP-SC1* failed to be induced by either *Lp^NC8^* or *Δdlt_op_* in *Dredd* larvae, both bacterial strains readily activated *PGRP-SC1* expression in control larvae with only minimal difference between the two bacterial strains (Fig. 4D). These results therefore indicate that the *Δdlt_op_* deletion only marginally alters the Dredd-dependent PG signal in the wild-type host, and the effect mediated through bacteria bearing D-alanylated TA in their cell wall likely activates another host commensal-sensing pathway in addition to the PGRP-LE/Imd/Relish cascade.

Next, we wished to examine if this second commensal sensing host pathway genetically interacts with the Dredd-dependent PG sensing pathway, and how such interaction affects expression of gut peptidases in response to *Lp.* We therefore analysed the expression of the intestinal peptidase, *Jon66Ci* and *Jon66Cii* in *Dredd* larvae monoassociated with either *Lp^NC8^* or *Δdlt_op_* strain. In contrast to control larvae, their expression in *Dredd* mutants was significantly reduced at all larval stages tested in the *Δdlt_op_* strain as compared to the *Lp^NC8^* strain (Fig. 4B-C). This result, combined with our expression study of *PGRP-SC1*, illustrates that the mechanism sensing bacteria bearing D-alanylated TA in their cell wall acts in concert with Dredd-dependent PG sensing to elicit optimal expression response from *Jon66Ci* and *Jon66Cii* loci. In the presence of *Δdlt_op_* mutation, the host mechanisms sensing bacteria bearing D-alanylated TA in their cell wall is impaired, and the *Dredd* mutation exacerbates the Δdlt_op_-related host response, which culminates to diminish intestinal peptidase expression. Since the *Δdlt_op_* mutation abolishes the D-alanine esterification of bacterial TA, we propose that the host enterocytes sense and signal the presence of *Lp* cells by at least two mechanisms, (1) through PGRP-LE-mediated PG fragment recognition and Imd/Relish signalling and (2) sensing of bacteria bearing D-alanylated TA in their cell wall and signalling by yet to discover host mechanisms.

Finally, we probed the consequence on *Lp* mediated larval growth promotion of altering both signals. We previously showed that altering Dredd-dependent PG sensing in the enterocytes was not sufficient to alter *Lp*-mediated *Drosophila* growth promotion (Erkosar et al., 2015). Here we compared the larval growth and maturation of control and *Dredd* animals upon association with either the *Lp^NC8^* or *Δdlt_op_* strains. We observed a significant effect on both larval growth (Fig. 4E) and maturation (Fig. 4F-G) of coupling the *Δdlt_op_* bacterial genotype and the host *Dredd* genotype. Taken together these results demonstrate that both bacterial PG and additional signals from bacteria bearing D-alanine esterification of TA in their cell wall, which are lacking in the *Δdlt_op_* strain, are required for optimal *Lp* mediated *Drosophila* growth and maturation upon chronic undernutrition.

## DISCUSSION

The results obtained in this study identify the *L. plantarum pbpX2-dltXABCD* operon as encoding an important bacterial functionality required to sustain a host-commensal dialogue that is beneficial to the host. Deletion of *pbpX2-dltXABCD* results in impairment of *Lactobacillus plantarum* (*Lp*) mediated *Drosophila* gut peptidases expression, larval growth and maturation. We show that *pbpX2-dltXABCD* is a key genetic determinant shaping *Lp* cell wall and cell shape via D-alanine esterification of TAs. Moreover, we reveal that these changes in the cell envelope architecture and composition are critical for bacterial sensing and signaling by *Drosophila* enterocytes, which underlies the beneficial interaction between *Drosophila* and its symbiont *Lp*.

Despite *Lp* being model lactic acid bacteria, *Lp* random transposon mutagenesis is difficult to achieve due to low-transformation efficiencies and/or instability of the transposon-delivering vector (Gury et al., 2004). Here, we successfully employed the P_junc_-TpaseIS_1223_ system (Licandro-Seraut et al., 2012) by constructing a random transposon mutant library in *Lp^NC8^.* We collected and screened 2091 transposon insertion mutants covering 1218 genes and tested around 42% of the predicted protein coding sequences in the *Lp^NC8^* genome (Axelsson et al., 2012). By increasing the number of transposon insertion mutants one could target virtually all non-essential genes. Upon screening of our library for its ability to promote larval growth after association with each one of the insertion mutants, we identified 7 insertions that severely impaired *Lp* growth promotion phenotype. Though significantly different from *Lp^NC8^*, none of the insertions reflects a complete loss-of-function phenotype that completely mimics the GF condition. This observation suggests that *Lp* growth promotion effect is probably multifactorial and to achieve such a “GF-like” phenotype we would need to target multiple bacterial functions.

Among the 7 mutated regions, we were particularly interested by the one impacting the *pbpX2-dltXABCD* operon, which is involved in *Lp* cell wall biogenesis and remodelling, and whose deletion impaired significantly *Drosophila* larval growth and interfered with host peptidase expression. Cell wall structure, composition and organization play major roles in host-bacteria dialogue as they represent the core bacteria components that trigger the initial microbe sensing response in the host. *pbpX2* encodes a putative D-D-carboxypeptidase with homology to various low-molecular weight penicillin binding proteins (Palumbo et al., 2006). The presence of *pbpX2* upstream of the *dltXABCD* genes is a unique feature of *Lp* genomes (Palumbo et al., 2006), but the physiological role of *pbpX2* and the importance of the genetic linkage between *pbpX2* and the *dltXABCD* genes remain unknown. The *dltXABCD* genes are well described in several gram-positive bacteria as being responsible for the esterification of teichoic acids with D-alanine (D-ala) (Grangette et al., 2005; Perego et al., 1995; Peschel et al., 2000). Teichoic acids (TA) are anionic polymers localized within the gram-positive bacteria cell wall, representing up to 50% of the cell envelope dry weight (Neuhaus and Baddiley, 2003). There are two types of TA: wall teichoic acids (WTA), which are covalently bound to peptidoglycan, and lipoteichoic acids (LTA), which are anchored on the cytoplasmic membrane (Neuhaus and Baddiley, 2003). The current model states that initially DltA ligates D-ala onto the carrier protein DltC. With the help of DltB, D-ala is then transferred from DltC to undecaprenyl phosphate crossing the membrane, where DltD transfers the D-ala to LTA (Reichmann et al., 2013). Although still not fully understood, it seems that D-alanyl-LTA serves as donor for D-ala substitutions in WTA (Reichmann et al., 2013). Despite being encoded upstream of *dltA* in several gram-positive bacteria, the role of DltX in TA D-alanylation remains unknown. *Lp^NC8^* carries in its genome the genetic information to produce both types of TA (Martino et al., 2016). Attachment of D-alanine substituents to these structures is an important mechanism by which bacteria modulates surface charge, whose level has a major impact on TAs functionalities, such as the control of autolysis (Palumbo et al., 2006), maintenance of cell wall morphology (Perea Vélez et al., 2007) and signalling with cells of their animal host (Grangette et al., 2005; Tabuchi et al., 2010). *Lp* strains lacking D-ala substitutions on their TAs reduce secretion of proinflammatory cytokines but better stimulate IL-10 production in peripheral blood monocytes when compared to its WT, showing the importance of TA D-alanylation in immunomodulating properties in mammals (Grangette et al., 2005; Rigaux et al., 2009). To our knowledge, direct sensing of TA and downstream signalling events have never been reported in *Drosophila*, yet D-alanylation of WTA seems an important bacterial feature impacting immunomodulation since its hampers recognition of *Staphylococcus aureus* PG by the pattern recognition receptor PGRP-SA (Tabuchi et al., 2010).

By quantification of D-alanine released from whole cells we determined that *pbpX2-dltXABCD* operon is responsible for D-alanylation of TA in *Lp^NC8^.* Moreover, the absence of the *pbpX2-dltXABCD* operon impacts cell morphology in *Lp^NC8^* while cell viability or growth dynamics are not affected in contrast to other lactobacilli strains (Palumbo et al., 2006; Perea Vélez et al., 2007). The consequences of deleting *dltXABCD* genes in lactobacilli are therefore context dependent. Cell morphological changes are frequently associated with defects in septum formation and modifications of autolysis activities. Therefore, our results suggest that PG remodelling might be affected, probably due to cell envelope charge modification leading to malfunction of cell wall hydrolases as previously reported in *L. rhamnosus^GG^* (Perea Vélez et al., 2007). This study highlights the role of *pbpX2-dltXABCD* and their impact on TA D-ala esterification for a bacterial beneficial effect on its host, a feature that is traditionally associated with pathogenesis and antibiotic resistance of gram-positive bacteria (Kristian et al., 2005; 2003; Peschel et al., 2000). Our results strengthen the role of TA modifications on commensal-host interactions and pave the way to further studies aiming at pinpointing the effect of *pbpX2-dltXABCD* function on *Lp* cell biology and physiology as well as probing the physiological role of *pbpX2.*

We previously established that the promotion of *Drosophila* linear growth by *Lp* is partly dependent on increased expression of host peptidases. We also demonstrated that such increased peptidases expression is partly controlled by PGRP-LE/Imd/Relish signalling cascade (Erkosar et al., 2015), a signalling pathway triggered by PG fragments (Leulier et al., 2003). Here we show that *pbpX2-dltXABCD* dependent bacterial signal(s) also contribute(s) to host intestinal peptidase induction and *Drosophila* larval growth. In this study, we observed in WT hosts a slightly reduced peptidases expression upon association with the *Δdlt_op_* strain when compared to the WT *Lp* strain. This differential peptidases expression is exacerbated in animals impaired in Imd signalling. Specifically, in *Dredd* mutants, peptidases expression upon *Δdlt_op_* strain association is close to the GF condition. Moreover, poor peptidases expression upon *Δdlt_op_* in *Dredd* mutants correlates to poor larval growth and maturation. Previously, we have established that the promotion of larval growth and maturation by *Lp* requires optimal expression of intestinal peptidases partially through PG-dependant Imd signalling. Imd deficient individuals have impaired PG sensing; yet in this context *Lp* still significantly promotes peptidases induction and larval growth. This observation indicates that *Lp* must be detected by additional sensing mechanisms beyond PG recognition. In the current study, we show that the absence of *pbpX2-dltXABCD* genes in *Lp^NC8^* depletes D-ala from TA, we therefore propose that bacteria with cell walls deprived of D-alanylated TA lack the additional bacterial signal sensed by *Drosophila* enterocytes to trigger intestinal peptidases induction and systemic growth (Fig. 5). At this stage, we propose that D-alanylated TA are directly sensed by *Drosophila* enterocytes but we cannot exclude the possibility that enterocytes directly sense other bacterial signals that would be altered in the *Δdlt_op_* context.

**Figure 5.**
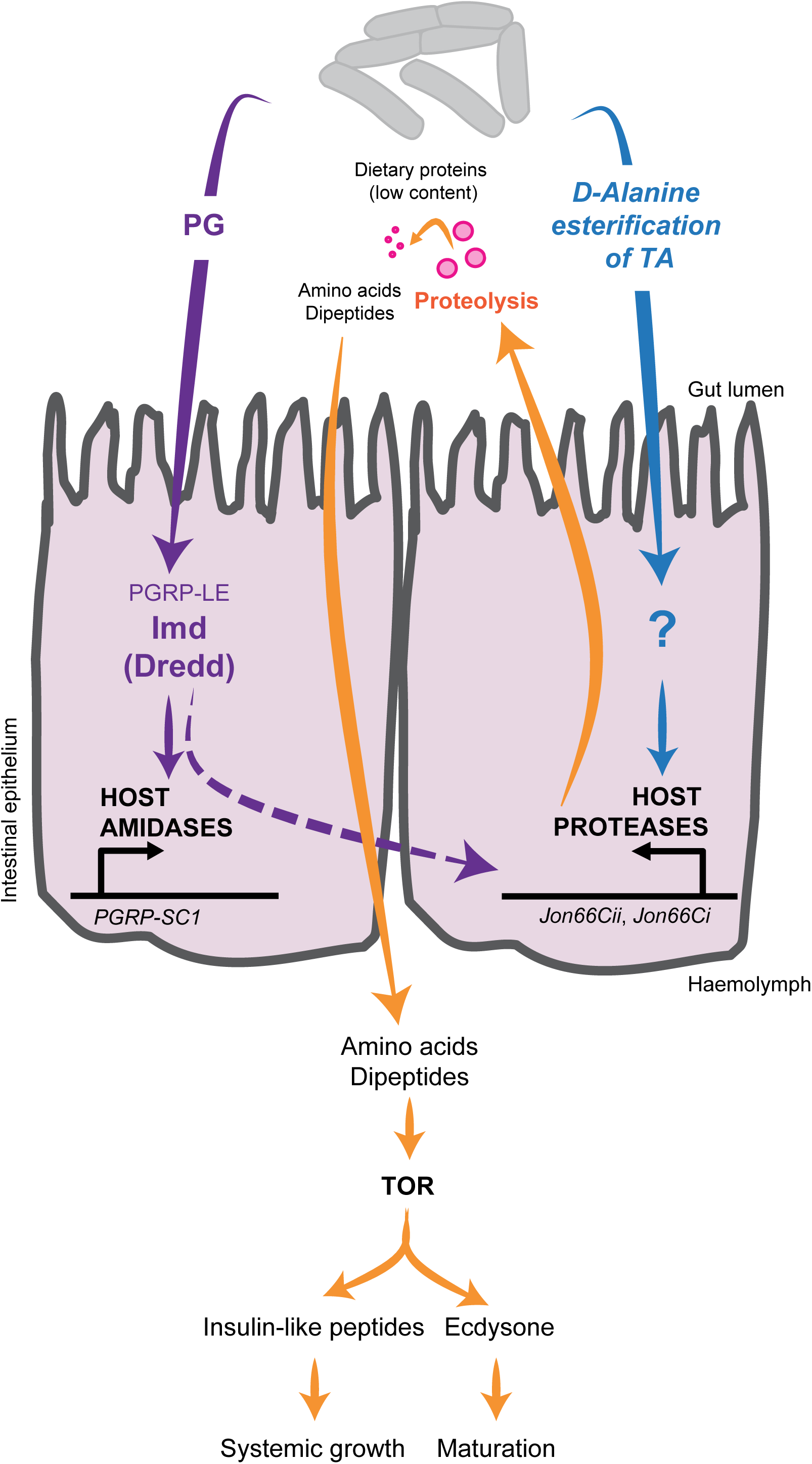
Cell envelope components of *L. plantarum* induce host intestinal digestive enzyme expression via a PG-responsible NF-κB-dependent signaling pathway and unknown signaling cascade(s), triggered by TA D-alanylation dependent signal. We previously established that detection of *L. plantarum* peptidoglycan via the PGRP-LE/Imd/Relish signalling cascade promotes, at least partly, peptidase expression in enterocytes. This leads to increased intestinal proteolytic activity, enhanced dietary protein digestion and improved amino-acid uptake in the host, conducting to increased TOR signalling activities in endocrine tissues and sustained production of systemic growth factors despite chronic undernutrition (Erkosar et al., 2015). Here we report that intestinal peptidase expression is increased upon recognition of both *Lp* PG and an additional signal dependent on esterification of teichoic acids with D-alanine.

Our study unravels a central molecular dialogue engaged between chronically undernourished *Drosophila* and its commensal partner *L. plantarum*, which supports the beneficial nature of their symbiosis. Given the recent demonstration of the conservation of the beneficial effects of *Lactobacillus plantarum* on the linear growth of chronically undernourished gnotobiotic mouse models (Schwarzer et al., 2016), our results therefore pave the way to probing whether this molecular dialogue is conserved in mammals.

## MATERIALS AND METHODS

### *Drosophila* diets, stocks and breeding

*Drosophila* stocks were cultured as described in (Erkosar et al., 2015). Briefly, flies were kept at 25°C with 12/12 hours dark/light cycles on a yeast/cornmeal medium containing 50 g/L of inactivated yeast. The poor yeast diet was obtained by reducing the amount of inactivated yeast to 6 g/L. Germ-free stocks were established as described in Erkosar et al. (2014). Axenicity was routinely tested by plating serial dilutions of animal lysates on nutrient agar plates. *Drosophila y,w* flies were used as the reference strain in this work. The following *Drosophila* line was also used: *y,w,Dredd^F64^* (Erkosar et al., 2015; Leulier et al., 2000).

### Bacterial strains and growth conditions

Strains used in this study are listed in Table S2. *E. coli* strains were grown at 37°C in LB medium with agitation. *L. plantarum* strains were grown in static conditions in MRS media at 37°C, unless differently stated. Erythromycin antibiotic was used at 5 μg/mL for *L. plantarum* and 150 μg/mL for *E. coli.*

### Random transposon mutagenesis of *L. plantarum^NC8^*

*L. plantarum* mutagenesis was performed using the P_junc_-TpaseIS_*1223*_ system as previously described by Licandro-Seraut et al. (2012) and Perpetuini et al. (2013). Briefly, electrocompetent *L. plantarum^NC8^* cells were first transformed with pVI129, resulting in the NC8pVI129 strain. Electrocompetent cells of NC8pVI129 strain were transformed with pVI110, plated on MRS plates supplemented with 5 μg/mL of erythromycin and incubated for 48 h at 42°C to select for integrants. 2091 tn-insertion mutants were individually stored at −80°C.

### Library high throughput insertion tracking: library construction and deep sequencing

Genomic DNA was extracted from each transposon insertion mutant by pools of 96 (UltraClean Microbial DNA isolation kit, MoBio). 22 DNA pools were quantified using Qubit system 2.0 (Invitrogen) and mixed together in equimolar proportion (4 μg / pool). DNA was sheared (1μg) following the manufacturer instructions using the S220 focused ultrasonicator (Covaris) to obtain a fragment distribution length from 100 bp to 1 kb, with an average peak around 400 bp. All the fragmented DNA material was then used to build a library using the Ion Xpress Plus gDNA Fragment Library Preparation kit (Thermofisher) following the protocol of the kit. However, only the P1 adapter was ligated at this step and not both adapters, A and P1, as usual. We designed a biotinylated fusion primer (named Primer Fusion A-IRL hereafter) containing: the sequence of the A adapter, the IRL sequence (transposon-specific primer IRL described in Scornec et al. (2014)) and a biotin at the 5’ end (reference Biot-TEG, Eurogentec) (Table S3). A PCR amplification was performed using the P1 primer and the primer fusion A-IRL, and the ligated-P1 DNA as a template. Reagents in the PCR mix were as follow: 1 μl of each primer at 10 μM, 47 μl of Platinum PCR SuperMix High Fidelity (Thermofisher), 1 μl of DNA at 50ng/μl for a final volume of 50 μl. After a denaturation step of 5 min at 95°C, 20 cycles of amplification were performed (15 s at 95°C; 15 s at 58°C; 1 min at 70°C), followed by a final extension step of 5 min at 70°C. The aim of this amplification step is to target the IRL sequence in the P1-library fragments, and add simultaneously the A adapter to build the final library, both A and P1 adapters being necessary for the subsequent sequencing. The fragments in the good configuration (meaning containing the biotinylated primer) were selected using Streptavidin magnetic beads (Dynabeads MyOne Streptavidin C1, Invitrogen) according to the manufacturer instructions. The biotinylated-selected amplicons were then purified using Qiagen MiniElute kit in order to eliminate the sodium hydroxyde, and re-amplified for 5 cycles following the classical re-amplification library protocol described in the Ion Xpress Plus gDNA Fragment Library Preparation kit (Thermofisher). The library was then size-selected using the E-gel Electrophoresis system (Invitrogen) in order to select fragments from 350 to 450 bp in length. The library was qualified according to the concentration and distribution profile using the TapeStation 2200 (Agilent). The diluted library (6 pM) was amplified through emulsion PCR using the Ion PGM Template OT2 400 kit (Thermofisher). Finally, the enriched library was loaded into a 314v2 chip and sequenced on the Ion PGM sequencer with the Ion PGM HiQ chemistry.

### Library high throughput insertion tracking by deep sequencing: bio-informatics analysis

A dedicated R script has been created to obtain mapping information of the reads on *Lp^NC8^* genome. The script takes in input the alignment sam file of the reads on the reference genome and compares the positions of the reads with an annotation table of the *Lp^NC8^* genome obtained with Geneious from the *Lp^NC8^* PubMed genbank file. It returns a table saying for each read if it maps a gene partially or totally or if it maps an intergenic region.

### BLAST Ring Image Generator (BRIG)

Visual analysis of the distribution of the transposon insertions was done by using BRIG (Alikhan et al., 2011; http://sourceforge.net/projects/brig/). *Lp^NC8^* was used as the reference genome. BRIG uses CGView (Stothard and Wishart, 2005) to render maps and is operated using a graphical user interface. The BRIG method uses the software BLASTALL v 2.2.25+ for the searches. The analysis was done with default settings.

### Library screening for loss of growth promotion phenotype

The screen was performed in the poor yeast diet. Each transposon insertion mutant (1×10^8^ CFUs) was used to independently inoculate 20 *y,w* germ-free eggs and incubated at 25°C for 8 days. The number of pupae 8 days after egg laying was scored for each of the 2091 tn-insertion mutants. Those values were converted in z-scores (where μ=15,18 n=2091 and σ =3,53).

### Sequence analysis and mapping of transposon insertion site

pVI110 locus of insertion were confirmed for the 8 loss-of-function candidates as described by Perpetuini et al. (2013), with the following modifications. Genomic DNA was digested sequentially with ClaI and BstBI restriction enzymes (NEB). Digested fragments were ligated using T4 DNA ligase (NEB) accordingly to manufacture’s instructions. Products of ligation were transformed into *E. coli* TG1 thermo-competent cells, in which circularized fragments containing the transposon behave as plasmids. Plasmids were isolated and sequenced (Genewiz) with the primers OLB221 and OLB215 (Table S3). Identification of transposon target sequences was performed with the BLAST software from the National Center for Biotechnology Information (NCBI).

### Construction of *ΔpbpX2* and *Δdlt_op_* deleted strains in *L. plantarum^NC8^* using pG+host9

Independent markerless deletions on *pbpX2* and *pbpX2* to *dltD (dlt* operon) genes were constructed through homology-based recombination with double-crossing over. Briefly, the 5′- and 3′-terminal regions of each region were PCR-amplified with Q5 High-Fidelity 2X Master Mix (NEB) from *L. plantarum* NC8 chromosomal DNA. Primers contained overlapping regions with pG+host9 (Maguin et al., 1996) to allow for Gibson Assembly. PCR amplifications were made using the following primers: OL118/OL119 and OL120/OL121 *(pbpX2)*, OL144/OL145 and OL146/OL147 *(dlt_op_)* listed in Table S3. The resulting plasmids (Table S2) obtained by Gibson Assembly (NEB) were transformed into NC8 electrocompetent cells and selected at the permissive temperature (28°C) on MRS plates supplemented with 5 μg/mL of erythromycin. Overnight cultures grown under the same conditions were diluted and shifted to the non-permissive temperature (42°C) in the presence of 5 μg/mL of erythromycin to select single crossover integrants. Plasmid excision by a second recombination event was promoted by growing integrants at the permissive temperature without erythromycin. Deletions were confirmed by PCR followed by sequencing with primers OL126/OL127 *(pbpX2)* and OL148/OL149 (*dlt_op_*).

### Larval size measurements

Axenic adults were put overnight in breeding cages to lay eggs on sterile poor yeast diet. Fresh axenic embryos were collected the next morning and seeded by pools of 40 on 55 mm petri dishes containing fly food. 1×10^8^ CFUs (unless otherwise specified) or PBS were then inoculated homogenously on the substrate and the eggs. Petri dishes are incubated at 25°C until larvae collection. *Drosophila* larvae, 7 days after inoculation, were collected and processed as described by Erkosar et al. (2015). Individual larval longitudinal length of individual larvae was quantified using ImageJ software (Schneider et al., 2012).

### Developmental timing determination

Axenic adults were put overnight in breeding cages to lay eggs on sterile poor yeast diet. Fresh axenic embryos were collected the next morning and seeded by pools of 40 in tubes containing fly food. 1×10^8^ CFUs of each strain (unless otherwise specified) or PBS were then inoculated homogenously on the substrate and the eggs and incubated at 25°C. Pupae emergence was scored everyday until all pupae have emerged. D50 was determined using D50App which is a Shiny app that calculates for the pupae emerged during a developmental experiment, the day when fifty percent of the pupae emerged. It takes as input a table with the number of pupae emerged every day for each condition and calculates with a local linear regression the day when fifty percent of the pupae emerged.

### Bacterial loads analysis

For larval loads, *y,w* axenic eggs were inoculated with 1×10^8^ CFU of each strain and incubated at 25°C until collection. Larvae were harvested from the nutritive substrate and surface-sterilized with a 30 s bath in 70% EtOH under agitation and rinsed in sterile water. Pools of five larvae were deposited in 1.5 mL microtubes containing 0,75-1 mm glass microbeads and 500 μL of PBS. To access bacterial CFU in the fly nutritional matrix, microtubes containing food were inoculated with 1×10^7^ CFUs of each strain, independently. Tubes were incubated at 25°C until being processed. For bacterial load quantification, 0.75-1 mm glass microbeads and 500 μL PBS was added directly into the microtubes. Samples were homogenized with the Precellys 24 tissue homogenizer (Bertin Technologies). Lysates dilutions (in PBS) were plated on MRS agar using the Easyspiral automatic plater (Intersciences). MRS agar plates were then incubated for 24 h at 37°C. Bacterial concentration was deduced from CFU counts on MRS agar plates using the automatic colony counter Scan1200 (Intersciences) and its counting software.

### Phase contrast and fluorescent imaging

Microscopy was performed on exponentially growing cells (OD600 = 0.3). Bacterial cultures (10 ml) were washed in PBS, centrifuged (3000g, 10 min) and resuspended in 150 mL of PBS. Bacterial cells were stained with FM4-64 (TermoFisher Scientific) at final concentration 0.1 mg/ml and incubated 5 min at room temperature in the dark. Slides were visualized with a Zeiss AxioObserver Z1 microscope fitted with an Orca-R2 C10600 charge-coupled device (CCD) camera (Hamamatsu) with a 100× NA 1.46 objective. Images were collected with axiovision (Carl Zeiss) and cell width at the half of the cell lenght (mid-width) was determined on phase contrast images by ImageJ using the MicrobeJ plug-in (Ducret et al., 2016).

### Quantification of D-alanine from teichoic acids by high-performance liquid chromatography (HPLC)

D-alanine esterified to teichoic acids was detected and quantified as described by Kovács et al. (2006). Briefly, D-alanine was released from whole heat-inactivated bacteria by mild alkaline hydrolysis with 0.1 N NaOH for 1h at 37°C. After neutralization, the extract was incubated with Marfey's reagent (1-fluoro-2,4-dinitrophenyl-5-L-alanine amide; Sigma). This reagent reacts with the optical isomers of amino acids to form diastereomeric *N*-aryl derivatives, which can be separated by HPLC. Separation of the amino acid derivatives was performed on a C_18_ reversed-phase column (Zorbax Eclipse Plus C18 RRHD 2.1×50mm 1.8μm Agilent) with an Agilent UHPLC 1290 system with a linear elution gradient of acetonitrile in 20 mM sodium acetate buffer (pH 4) as described previously (Kovács et al., 2006). The eluted compounds were detected by UV absorbance at 340 nm. Quantification was achieved by comparison with D-alanine standards in the range of 100 to 1500 pmol. Mean values were obtained from five independent cultures with two injections for each.

### RNA extraction and RT-qPCR analysis

Axenic *y,w* and *y,w,Dredd* eggs were inoculated with 1×10^8^ CFU of *Lp^NC8^* and *Δdlt_op_* strains independently or kept axenic. Larvae were size matched for the three conditions and harvested at two different sizes: ≈2,5 mm and ≈4 mm. RNA extraction of five replicates of ten larvae for each condition was performed as described by Erkosar et al. (2015). RT-qPCR was performed using gene-specific primer sets (Table S3) as described by Erkosar et al. (2015). Results are presented as the value of ΔCt^gene^/ΔCt^rp49^.

### Statistical analysis

Data representation and analysis was performed using Graphpad PRISM 6 software (www.graphpad.com). Mann Whitney’s test was applied to perform pairwise statistical analyses between conditions. Student’s T test with Welch correction was performed to determine the significance of differences in gene expression levels between *Lp^NC8^* and *Δdlt_op_* inoculated samples. Kruskal Wallis test was applied to perform statistical analyses between multiple (n>2) conditions.

## Acknowledgments

The authors would like to thank Maura Strigini, Gilles Storelli and Dali Ma for critical reading and editing of the manuscript, Christophe Grangeasse for helpful discussions and help with bacterial cells imaging, the Arthro-Tools and PLATIM platforms of the SFR Biosciences (UMS3444/US8) for providing *Drosophila* and imaging facilities, the IGFL sequencing platform for deep sequencing, Pascale Serror for pG+host9 and Hélène Licandro-Seraut for P_junc_-TpaseIS_*1223*_, system. RCM thanks the “Fondation pour la Recherche Médicale” for financial support through a postdoctoral scholarship SPF20140129318. This work was funded by an ERC starting grant (FP7/2007-2013-N°309704). FL lab is supported by the FINOVI foundation, the “Fondation Schlumberger pour l’Education et la Recherche” and the EMBO Young Investigator Program. The authors declare no conflict of interests.

## Author contributions

FL supervised the work. RCM and FL designed the experiments. RCM and HG performed the experiments. MS performed bacterial cell imaging. BG and SH designed and performed high-throughput insertion tracking by deep sequencing. MEM and PJ performed the insertions sites bioinformatics analysis. PC performed D-alanine quantifications. RCM, PC, MPCC, MS, and FL analyzed the results. RCM and FL wrote the paper.

## Supplementary materials

**Figure S1.**
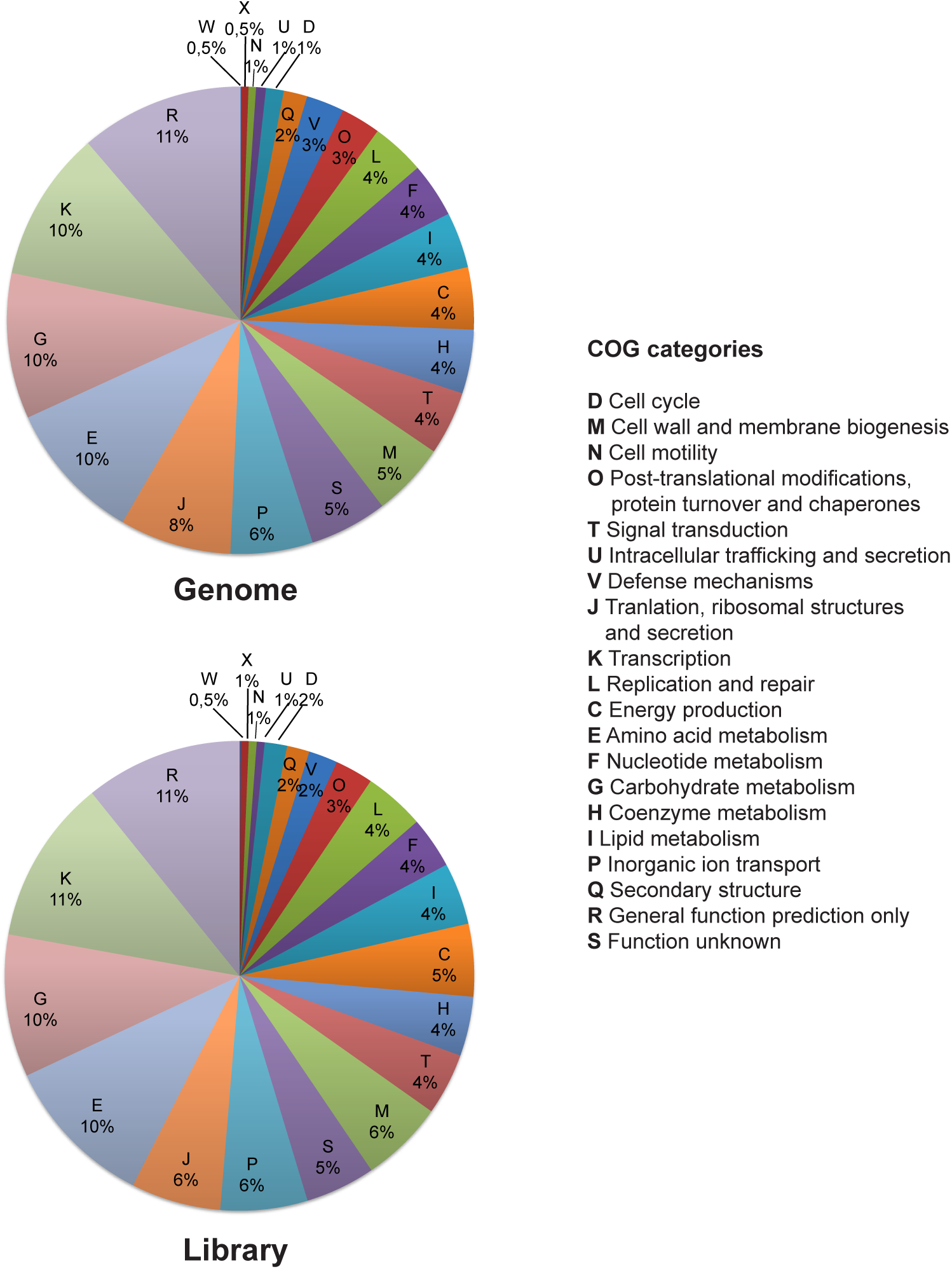
Analysis of transposon insertions in the genome of 2091 *L. plantarum^NC8^* mutants: Relative abundance of clusters of orthologous groups (COG) functional categories of genes in *Lp^NC8^* genome and in the transposon library. COG categories are indicated on the right side of the panel.

**Figure S2.**
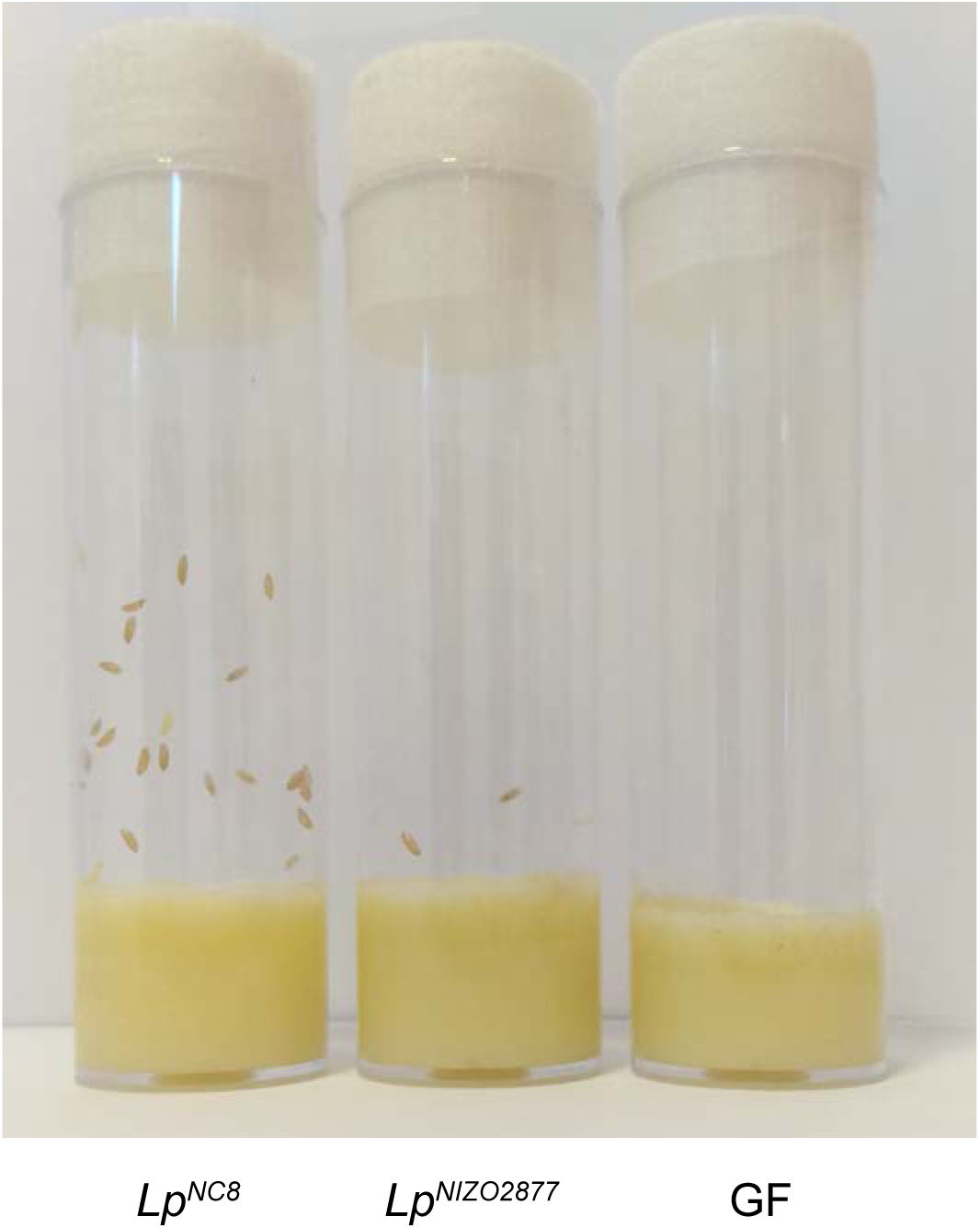
Representative images of developmental timing tubes with germ-free eggs inoculated with a strong growth promoting strain *(Lp*^NC8^), a mild growth promoting strain *(Lp^NIZO2877^*) and PBS (for the GF condition), 8 days after association.

**Figure S3.**
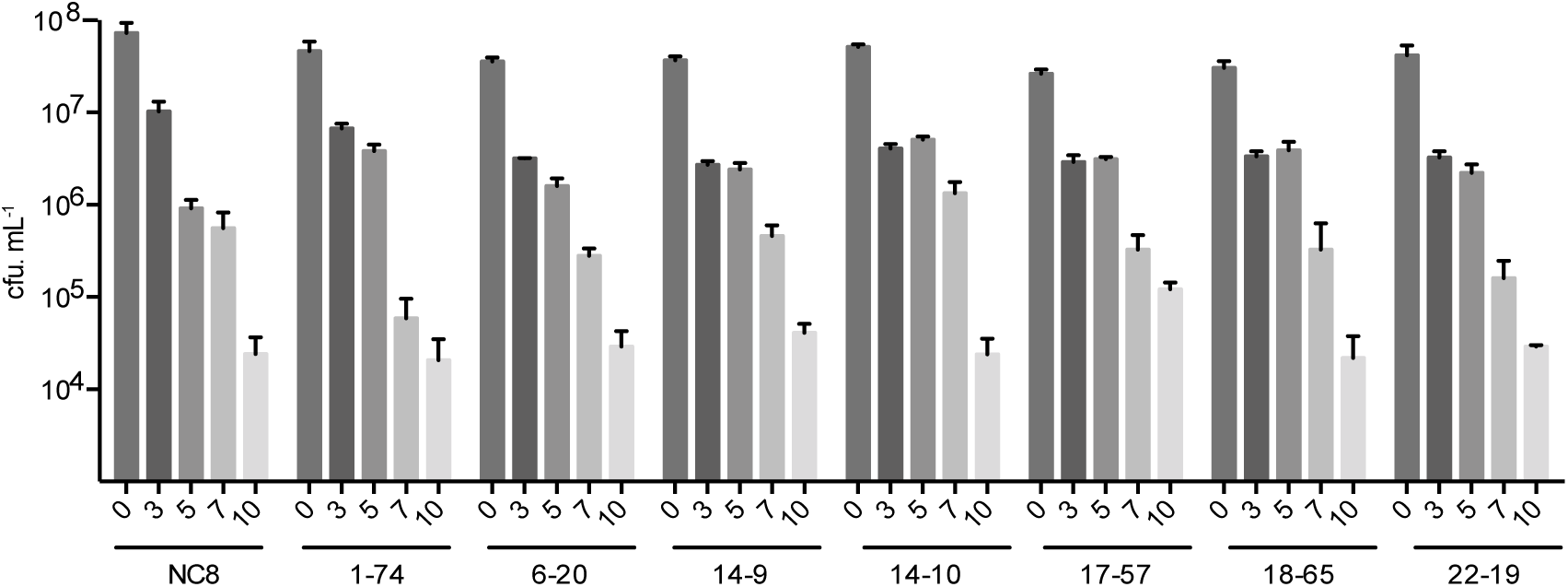
Evolution of the number of CFUs on fly food at days 3, 5, 7 and 10 after inoculation for the 7 candidates selected from the secondary screen.

**Figure S4.**
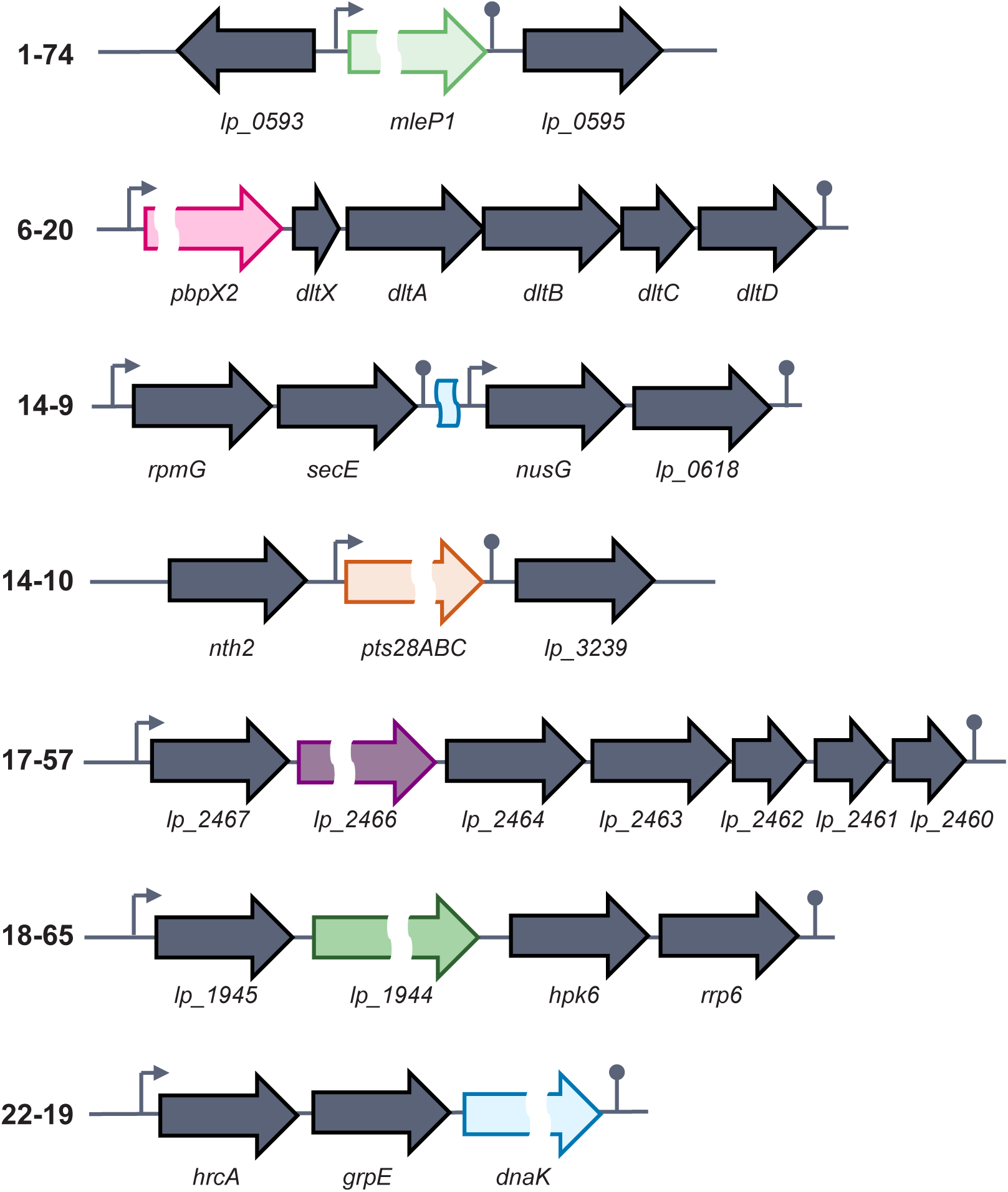
Genes disrupted by the transposon in the loss-of-function candidates (genomic context and operon predictions made with fgenesB (Solovyev and Salamov 2011). Genes and intergenic region hit by the transposon are highlighted in color. Arrows represent the predicted operon promoters and loops represent their terminators.

**Table S1.** Transposon insertions in coding regions. Gene name, position in *Lp^NC8^* genome (contig number and position of the insertion within the contig), orthologous gene (OG), locus tag in *Lp^WCFS1^* genome, COG and category.

**Table S2.**
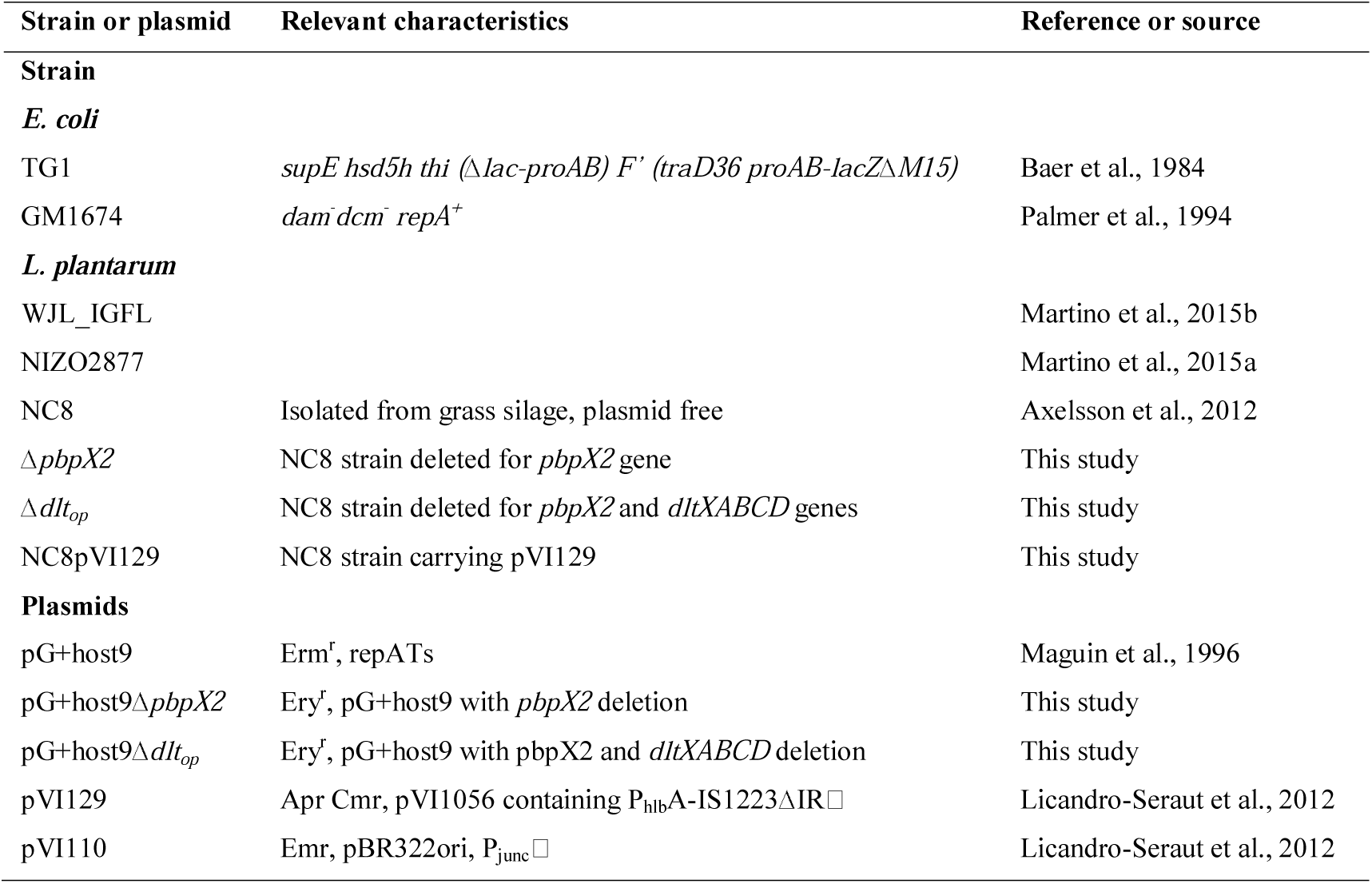
Bacterial strains and plasmids used in this study.

**Table S3.**
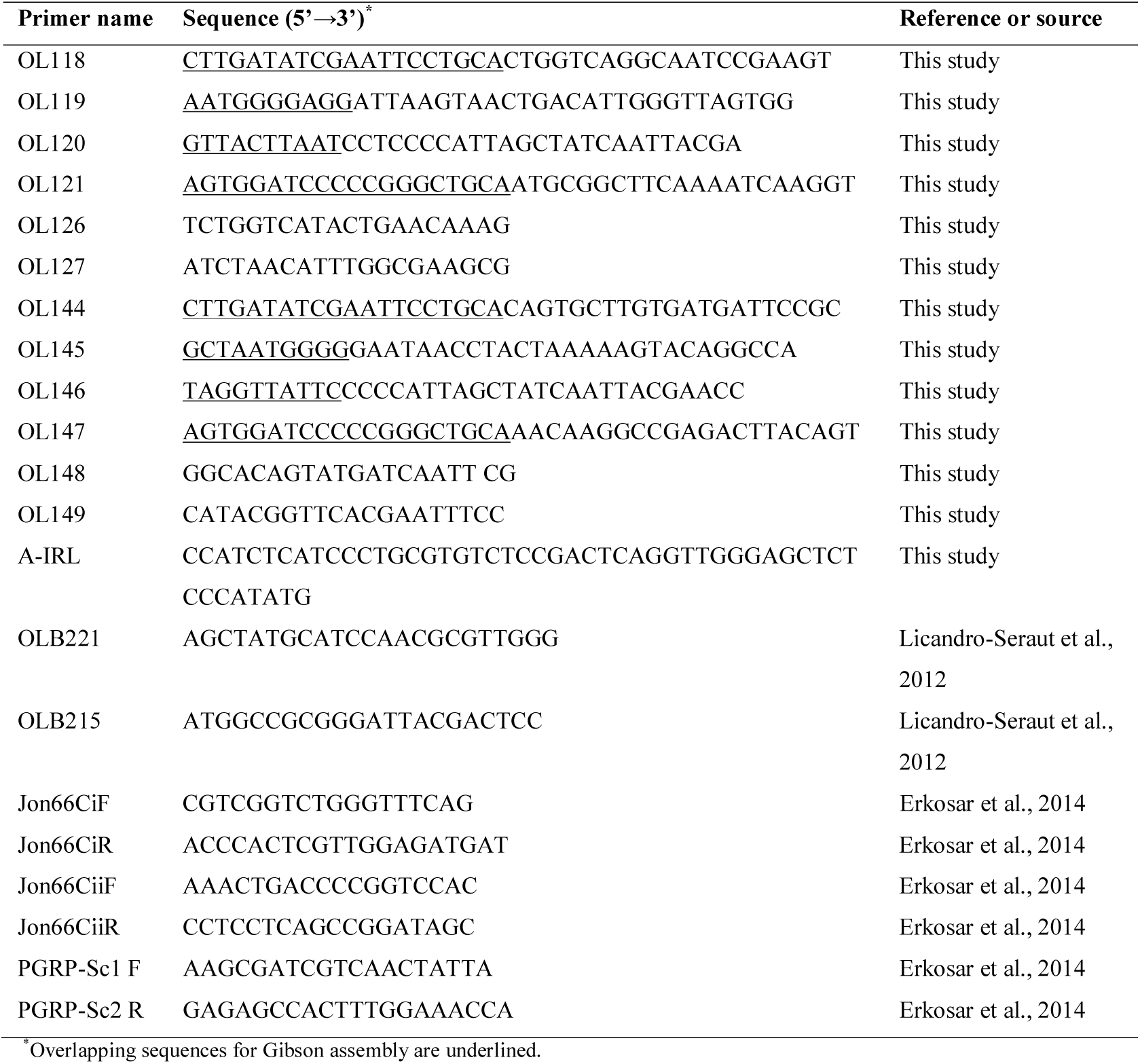
Primers used in this study.

